# GeneCAD: Plant Genome Annotation with a DNA Foundation Model

**DOI:** 10.1101/2025.10.31.685877

**Authors:** Zong-Yan Liu, Ana Berthel, Eric Czech, Edgar Marroquin, Michelle Stitzer, Sheng-Kai Hsu, Matt Pennell, Edward S. Buckler, Jingjing Zhai

## Abstract

Accurate genome annotation is fundamental to biological discovery, yet identifying gene structures directly from DNA sequence remains a major challenge in complex genomes. We introduce GeneCAD, a sequence-only framework that predicts biologically coherent gene models without requiring species-matched transcriptomic or proteomic evidence. GeneCAD integrates lineage-specific DNA representations from the PlantCAD2 foundation model with a transformer encoder and a chromosome-scale conditional random field (CRF) to enforce structural constraints, such as splice-phase and feature order. To ensure high-quality supervision, we implement a curation strategy using a sequence-based masked-motif score to filter reference transcripts. As a primary validation across diverse angiosperms, including a complex allotetraploid, GeneCAD improves transcript F1 by approximately 9% over current tools like Helixer and BRAKER3, while sharpening boundary precision and achieving a best-in-class recovery of 86% of classical coding sequences. Furthermore, we demonstrate the framework’s modularity by adapting it to animal lineages through the substitution of the underlying DNA foundation model. While the long introns of vertebrates challenge full transcript reconstruction, the model remains highly effective at identifying individual exons. By connecting evolutionary signals with structured decoding, GeneCAD provides a versatile and scalable solution for high-fidelity genome annotation across the Tree of Life.

**Graphical Abstract:** 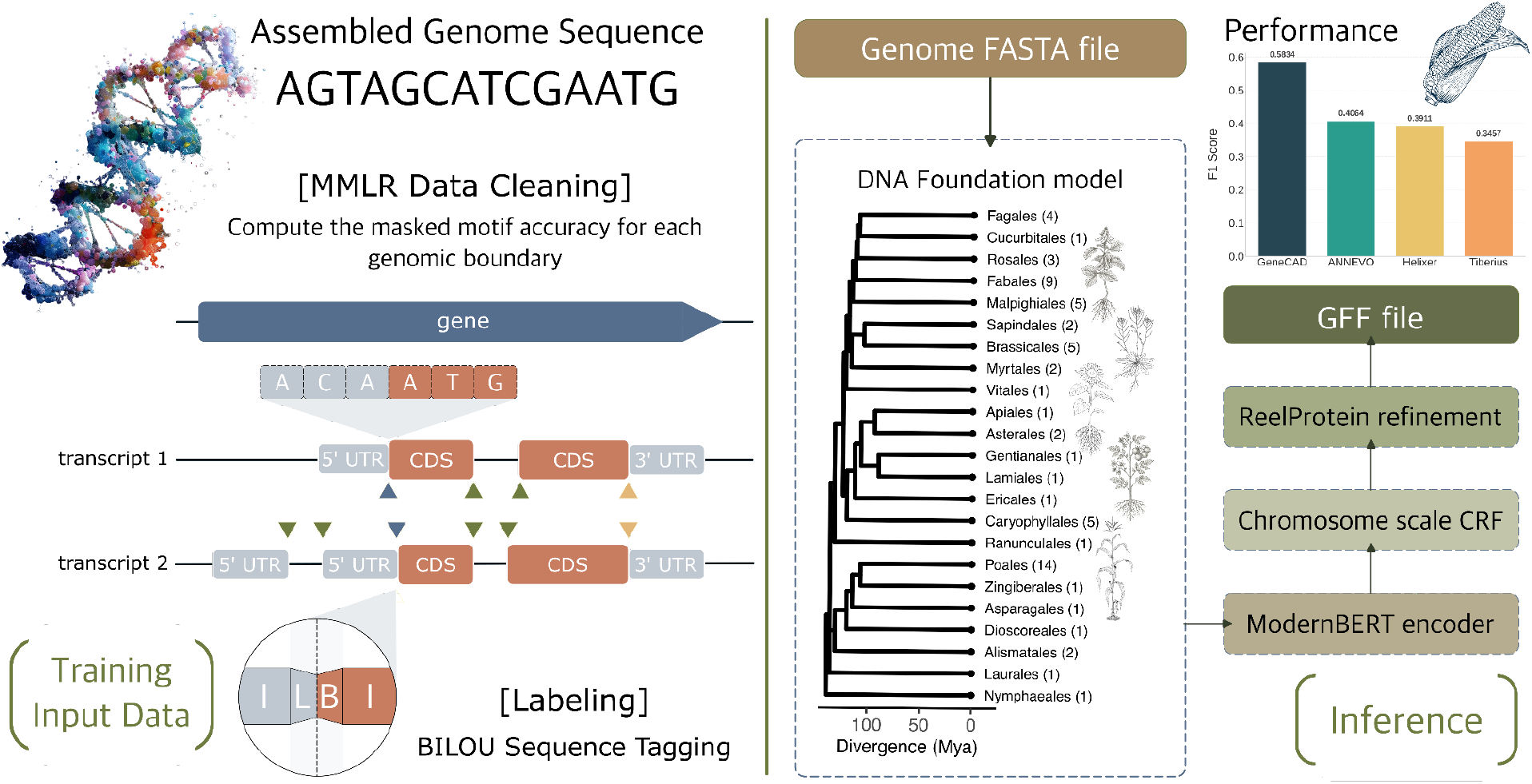

**Lay Summary:** Identifying where genes are located within a genome is a major challenge in biology, especially in plants with large and repetitive DNA. Current methods often rely on expensive laboratory data or struggle to find genes in “noisy” regions. We developed GeneCAD, a deep-learning tool that identifies genes using only the raw DNA sequence. By using AI models that recognize patterns shaped by millions of years of evolution, GeneCAD predicts gene structures with high accuracy and consistency. Our tests show that GeneCAD outperforms existing tools in plants and can be easily adapted for use in other species, including animals. By removing the need for costly lab experiments, GeneCAD provides an affordable way for researchers to quickly map the genomes of everything from rare wild plants to essential food crops.

## Introduction

Coordinated long-read consortia, most notably the Earth BioGenome Project [1], have generated thousands of chromosome-scale reference genomes across plants and animals and are scaling toward tens of thousands more. Realizing the full value of these assemblies now hinges on precise annotation, which enables gene function prediction, anchors genotype–phenotype associations, and accelerates advances from basic biology to broad genomic improvement. For newly assembled genomes, pipelines such as BRAKER3 [2] integrate species-matched RNA-seq and homologous protein evidence to train generalized hidden Markov model (HMM) based gene finders (e.g., GeneMark-ETP [3] and AUGUSTUS [4]) and to reconcile predictions (e.g., with TSEBRA [5]) for genome-wide gene calling. Homology detection, efficiently implemented with profile HMMs, provides a rapid and broadly applicable signal that complements the tissue- and stage-diverse transcriptomic and proteomic data required for optimal accuracy. However, RNA-seq can inflate false positives, including transposable-element (TE) fragments and pseudogenes [6], and substantial model inconsistency persists even in well-curated species like maize [7,8]. Proteome offers a stronger discriminator than RNA-Seq driven workflows, flagging about 28% of maize transcripts [9] as incorrectly annotated or nonfunctional while agreeing with classical and proteome-supported sets. Although proteomics yields high-confidence coding evidence, it remains expensive and low throughput, and many specimens provide only DNA with no accessible RNA, let alone proteome. Limited coverage therefore leads to false negatives, including missed exons and low-abundance isoforms [10]. For wild or rare plant specimens, generating comprehensive RNA-seq and proteomes is often infeasible and sequences are frequently fragmented. In the absence of these inputs, RNA- and protein-dependent pipelines can fail or return incomplete models, and homology support weakens with phylogenetic distance. Under such constraints, limited assay coverage, polyploidy, and repeat-rich genomes, which are particularly prevalent in the plant kingdom, further hinder genome annotation.

To reduce dependence on experimental data, sequence-only deep learning tools such as Helixer [11], Tiberius [12], and ANNEVO [13] predict genes directly from DNA without additional transcriptomic or homology inputs. These approaches are attractive because they scale across species and circumvent the need for assay generation. Because supervised models require large labeled datasets to achieve cross-species performance, they are typically trained on public reference annotations from public databases such as Phytozome [14], Ensembl [15], and RefSeq [16]. However, these resources vary in quality and consistency and contain known misannotations, including pseudogenes, TE-derived open reading frames, fused or split genes, and junctions reflecting pre-spliced RNA, which can propagate label noise during training. These issues are especially acute for functionally critical but highly duplicated, rapidly evolving immune-receptor families such as nucleotide-binding leucine-rich repeats (NLRs) [17], which often occur in repeat-dense clusters and are frequently misannotated in automated genomes. We therefore hypothesize that stronger curation of training data and structure-aware decoding are necessary to achieve consistent, biologically coherent performance across evolutionarily diverse lineages.

Recent advances in self-supervised foundation models have opened new possibilities for genome annotation at scale. Trained on raw DNA or protein, these models learn conservation-aware representations that transfer across species. Independent studies have likewise trained large nucleotide-level segmentation models on extensive annotations and reported strong transfer. For example, SegmentNT [18] built on the Nucleotide Transformer [19]. For DNA, representative models include PlantCAD2 [20], GPN-MSA [21], and Evo2 [22]; for proteins, ProtTrans [23], ESM3 [24]. PlantCAD2 was pretrained on 65 diverse plant genomes with an 8,192 bp context window, capturing long-range signals of evolutionary constraint across angiosperms, after which task-specific heads incorporate structural and functional priors. Using PlantCAD2, we demonstrate cross-species generalization with limited labels. However, nucleotide-level labeling alone does not constitute gene prediction; without structure-aware decoding, outputs can violate annotation rules and cannot be assembled into valid transcript models.

We introduce GeneCAD, a modular and lineage-agnostic framework for plant genome annotation that works directly from DNA. GeneCAD couples single-nucleotide-resolution representations from a lineage-appropriate foundation model, such as PlantCAD2 [20] for plant genomes, with a ModernBERT [25] head and a chromosome-scale conditional random field (CRF) [26] that enforces gene-structure constraints. Single-nucleotide BILOU tags [27] capture boundaries for UTRs, introns, and coding segments, and the CRF converts those scores into coherent transcripts that respect feature order and splice rules. To limit label noise, we curated a quality-aware curation strategy with a zero-shot masked-motif score and trained GeneCAD on a compact, phylogenetically diverse subset. A protein language model screen removes phase-consistent but implausible open reading frames. While GeneCAD achieves state-of-the-art performance in plants, we also tested the framework’s portability by adapting it to animal lineages using the Caduceus foundation model [28]. In these vertebrate tests, the model successfully identified individual exons but struggled to reconstruct complete transcripts due to the extreme length of animal introns. These results demonstrate that while the GeneCAD architecture is lineage-agnostic, transcript-level resolution in animals remains a distinct challenge for sequence-only models. Together, our results show that GeneCAD provides a robust, scalable solution for high-fidelity plant genome annotation and establishes a foundation for cross-lineage gene prediction.

## Materials and Methods

### Source Data and Lineage-specific Scaffolding

GeneCAD utilizes lineage-specific foundation models and reference datasets tailored to the target clades. For the primary plant validation, we began with the 65 plant genomes used to pretrain PlantCAD2 and applied a sequence-only screen to prioritize references for fine-tuning. For each annotated transcript, we queried PlantCAD2 [20] in a zero-shot setting to obtain recovery probabilities at the translation start codon, the translation stop codon, and the splice donor and acceptor dinucleotides. We combined these motif-level probabilities into a transcript-level masked-motif logistic regression score (MMLR) The logistic model was trained with positive-unlabeled learning, using classical *Zea mays* genes as positives and all other transcripts as unlabeled (Supplementary Note 1). We ranked species by mean MMLR and also considered assembly contiguity, BUSCO v5.4.3 completeness [37], and whether annotations were produced by evidence-supported pipelines that yield consistent models. This process selected five references: *Arabidopsis thaliana, Glycine max, Hordeum vulgare, Oryza sativa*, and *Populus trichocarpa* (Supplementary Table 1). To evaluate cross-lineage generalizability, we trained the framework on genomes from Homo sapiens (GRCh38), Mus musculus (GRCm39), Danio rerio (GRCz11), Xenopus tropicalis (v10), and Callithrix jacchus (mCalJac1.pat.X). The Gallus gallus (GRCg7b) genome was reserved exclusively for testing. Utilizing Ensembl-sourced assemblies, we integrated the Caduceus foundation model while maintaining the original modular architecture and task-specific decoding heads. MMLR was computed in a zero-shot manner and was used only for species and transcript selection. For the external check of the plant models, we assigned proteome support in *A. thaliana* [38], *Z. mays* [9,38], *S. lycopersicum* [39], and *O. sativa* [40] from public peptide evidence (Fig. 2b). A transcript was considered supported when at least one peptide uniquely mapped to its encoded protein sequence from the reference FASTA. These labels were used only for external check and were not used for GeneCAD training.

### Training Data Generation and Curation

Within each selected species, we retained transcripts with an MMLR score greater than 0.5. At loci with multiple isoforms, we designated the highest-scoring isoform as canonical. We standardized GFF3 files to harmonize feature hierarchies. We excluded transcripts that contained internal stop codons, non-canonical splice forms, or same strand overlaps with neighboring genes. Retained models were converted to a tabular schema of genes, mRNAs, exons, coding segments, and UTRs, and were paired with genomic coordinates to yield a distilled, non-redundant corpus for fine-tuning and validation (Fig. 2b,c). MMLR was computed as described above and in Supplementary Note 1. Per-species mean MMLR and retention counts are reported in Supplementary Table 1. This curation pipeline was applied identically to both plant and animal reference annotations to ensure a high-confidence ground truth for training.

### Independent Test Set for Rigorous Evaluation

To assess generalization across diverse evolutionary histories and genome architectures, we assembled two independent test sets that were strictly excluded from training. The primary plant test set consisted of five angiosperm species: *Juglans regia* (walnut), *Coffea arabica* (coffee), *Phaseolus vulgaris* (common bean), *Nicotiana sylvestris* (diploid tobacco), and *Nicotiana tabacum* (allotetraploid tobacco) [41–45]. These species span major angiosperm clades and include a diploid–allotetraploid pair, enabling evaluation across ploidy levels. Reference annotations were used as released and were not re-filtered with MMLR. Each assembly is supported by extensive experimental evidence, including RNA-seq, long-read Iso-Seq, and proteomics. Approximate separations from the nearest training lineage range from tens to more than 100 million years (Supplementary Table 1). To evaluate the model’s behavior on a different vertebrate lineage, we used the chicken (Gallus gallus, GRCg7b) genome as an independent animal test case. This species was reserved exclusively for testing to measure how well the framework translates to a genome not seen during training. We also evaluated *Zea mays* as an additional held-out case. *Z. mays* was not used to fine-tune GeneCAD and provides a high-evidence reference that is informative for model behavior. We used maize to quantify absolute performance and to analyze the effects of pretraining and curated training diversity, including ablations (Fig. 4). To avoid circularity, we did not include maize in the primary cross-tool benchmark because it appears in the training data of some comparator pipelines.

### Data representation and labeling

To ensure a unified processing pipeline across different lineages, we implemented a standardized data representation scheme compatible with both PlantCAD2 and Caduceus. Genomic FASTA files for all curated training and independent test sets were tokenized to integer IDs with a single-base vocabulary (A, C, G, T, N). Forward and reverse-complement token streams were stored in Zarr format to allow efficient random access during training and inference.

Per-base labels were derived from curated canonical transcripts. Four feature types were annotated at nucleotide resolution: CDS, intron, 5′ UTR and 3′ UTR. Intergenic sequence was assigned to a single Outside (O) class. For each contiguous span of a feature, we used the BILU scheme: the first base was tagged B, internal bases I, the last base L, and single-base spans U. These tags were applied separately for CDS, intron, 5′ UTR and 3′ UTR; all remaining bases were labeled Outside. In total, the labeling scheme defines 17 classes: four feature types, each with four BILU tags, and one Outside class [27].

### GeneCAD Model Architecture

#### Input Representation

GeneCAD uses a modular input layer that can incorporate representations from different DNA foundation models. Training sequences were sampled as 8,192 bp windows with a 4,096 bp step size, corresponding to 50% overlap. At each position, GeneCAD combines the contextual hidden state from the selected base encoder with a trainable 256-dimensional token embedding representing the local nucleotide token. When required, base-encoder hidden states were linearly projected so that, after concatenation with the token embedding, the per-base representation matched the head encoder dimension. In the configuration used here, this produced a 1,024-dimensional per-base representation (Fig. 2b). The base encoder, projection layer, token embeddings, ModernBERT head encoder, and classifier were optimized jointly during training, allowing pretrained representations and task-specific layers to adapt to gene-structure annotation. The resulting sequence representations were processed by an 8-layer ModernBERT head encoder, followed by the token-level prediction head and downstream decoding procedure (Fig. 1b).

**Figure 1.**
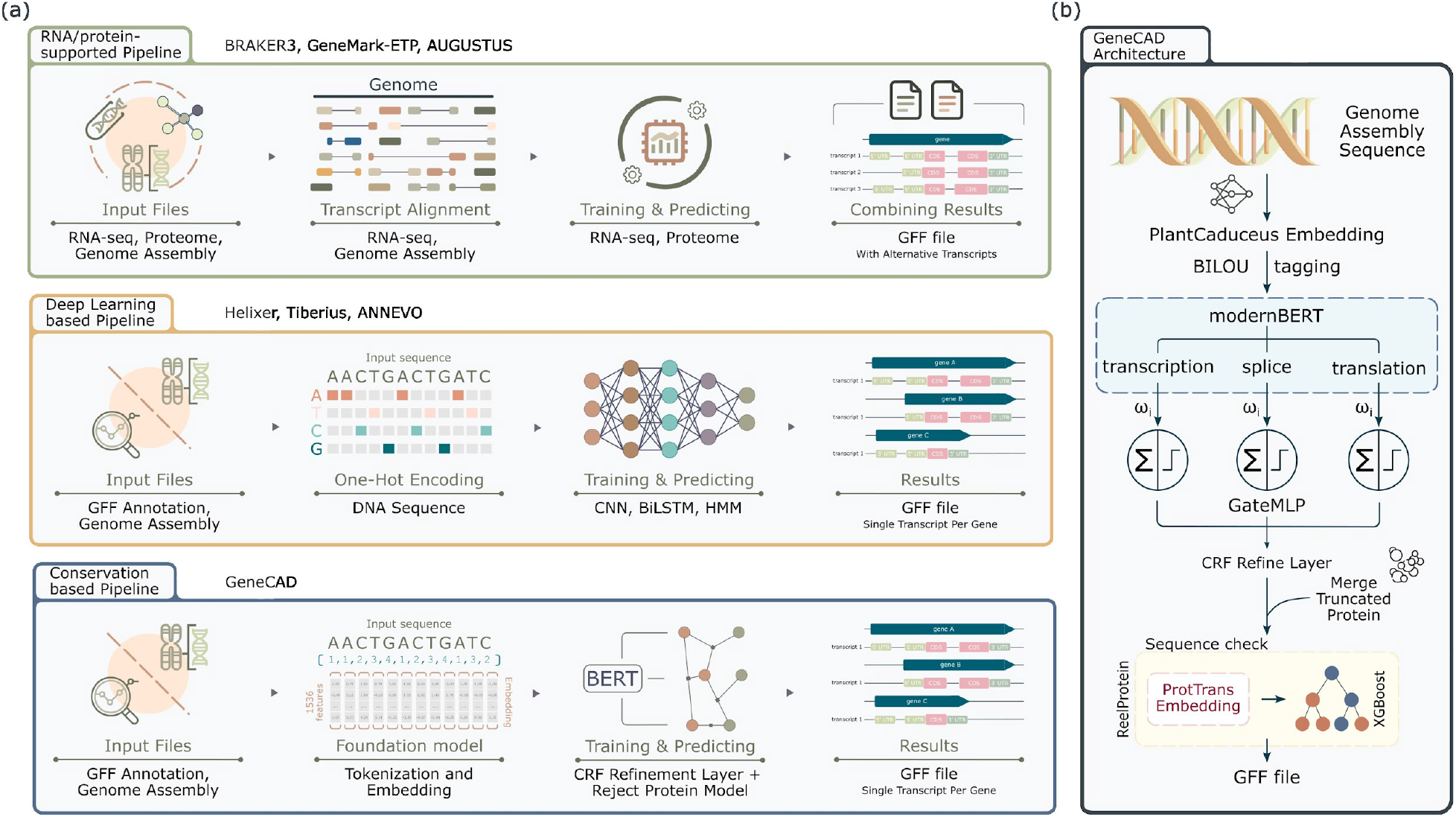
GeneCAD architecture compared with existing annotation pipelines. **(a) Three pipeline families**. Evidence-supported tools (for example BRAKER3/AUGUSTUS) align RNA-seq and proteomic data to produce multi-isoform annotations. Ab initio deep-learning tools (for example Helixer, Tiberius) operate only on DNA and often blur feature boundaries. GeneCAD uses a conservation-aware, foundation-model strategy: it ingests lineage-specific DNA embeddings and applies structured decoding to output one canonical, structurally coherent transcript per locus. **(b) GeneCAD architecture**. Genome sequence is embedded with a foundation model (here, PlantCAD2) and labeled at single-nucleotide resolution with BILOU tags. An eight-layer ModernBERT encoder produces contextual states that feed three task streams for transcription boundaries, splicing, and translation. A gated MLP fuses the streams to yield per-base logits. A chromosome-wide CRF with empirically derived transition constraints performs Viterbi decoding, enforcing valid feature order and splice consistency. Post-processing reconnects window-split loci when a single in-frame ORF is supported, and predicted CDSs are screened with ReelProtein. The final output is a GFF3 with one canonical transcript per gene.

**Figure 2.**
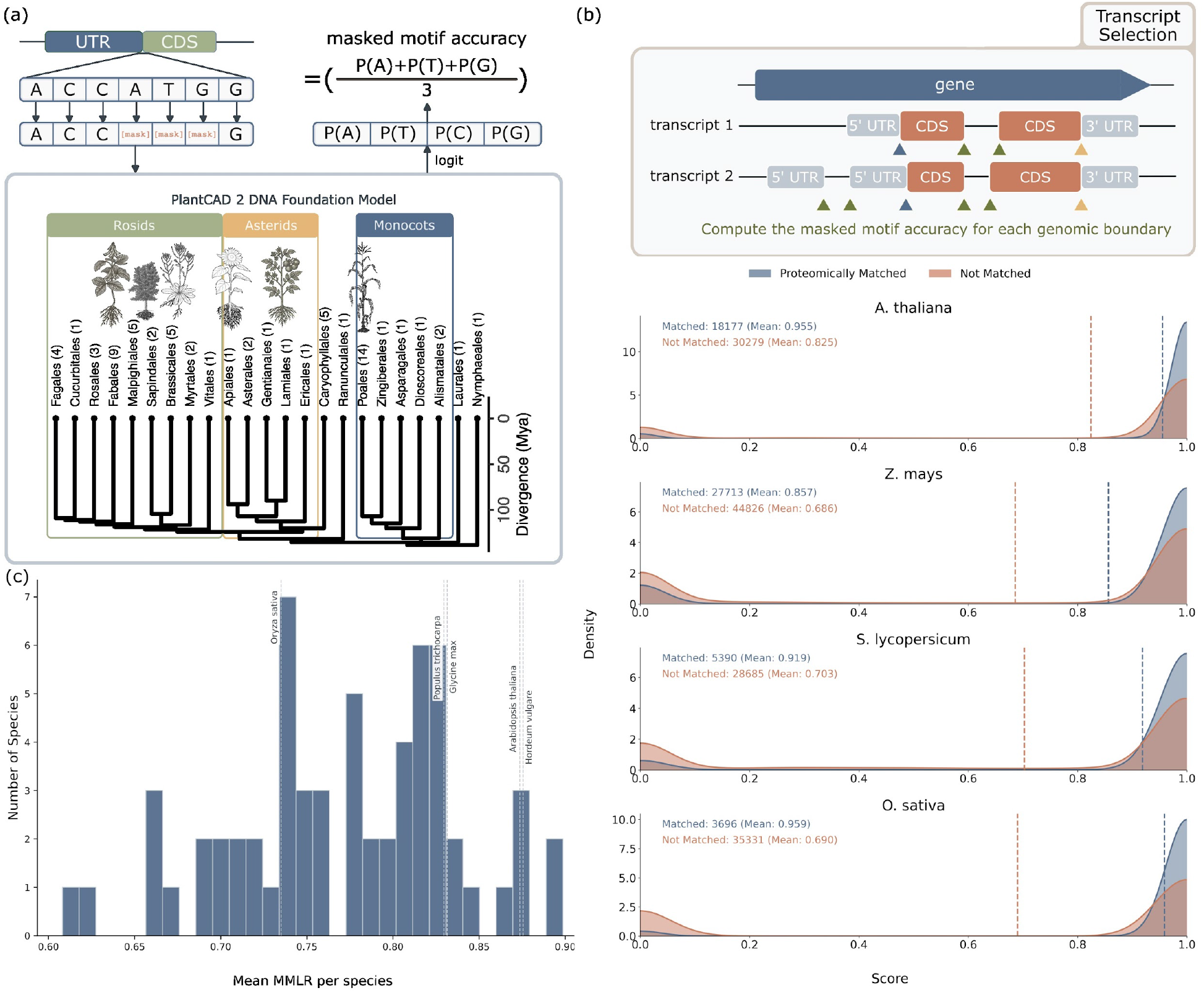
Masked motif logistic regression (MMLR) enables quality assessment and curation of reference genome annotations. **(a) Masked-motif scheme**. At each annotated boundary, translation start or stop and splice donor or acceptor, the focal nucleotides are masked. The foundation model (PlantCAD2) is queried zero-shot to recover the masked base from surrounding context. Per-motif probabilities are combined into a transcript-level masked-motif logistic regression score (MMLR). **(b) Transcript-level scoring and proteome support**. For each transcript, masking is applied at all coding and splice boundaries to yield one MMLR value used for selection. Density plots compare MMLR distributions for peptide-supported (blue) and unsupported (red) transcripts in four species (*Arabidopsis thaliana, Zea mays, Solanum lycopersicum, Oryza sativa*). Vertical dashed lines mark group means; the curation threshold is indicated at right in each panel. **(c) Species-level variation and reference selection**. Mean MMLR per species across the primary validation pretraining genomes shows broad variation. Dashed markers denote the five references selected for fine-tuning based on MMLR, assembly quality, and BUSCO completeness (*A. thaliana, Glycine max, Hordeum vulgare, O. sativa, Populus trichocarpa*).

#### ModernBERT Encoder

The head encoder was implemented as an 8-layer ModernBERT stack with a hidden size of 1,024 and eight attention heads per layer. It followed the standard ModernBERT layer design, with multi-head self-attention and a feed-forward block whose intermediate width was four times the hidden size [46]. Attention and MLP dropout rates were set to 0.1. The encoder preserved sequence length and returned a contextual state for every genomic position. These states were passed to a token-level classifier to produce per-base token logits (Fig. 2b).

#### Gated MLP Classification Head

Final per-base classification was performed with a gated MLP whose hidden width matched the feed-forward width of the ModernBERT head encoder. For the 1,024-dimensional configuration, the MLP used a 4,096-dimensional intermediate representation. At each position, a linear projection produced separate signal and gate vectors. The signal vector was activated with GELU and multiplied element-wise by the gate. The gated representation was then passed through dropout with a rate of 0.1 and linearly mapped to 17 token classes. These classes represented intron, 5′ UTR, CDS and 3′ UTR using BILU tags, together with a shared intergenic/background class.

#### Model Training and Optimization

Models were implemented in PyTorch with PyTorch Lightning and trained with bfloat16 mixed precision. We used AdamW with a learning rate of 2 × 10^−4^, by default AdamW momentum parameters, *β*_1_ = 0.9, *β*_2_ = 0.999, ε = 1 × 10^−8^, and a weight decay of 0.01. When the base encoder was trainable, its learning rate was set to one-tenth of the head learning rate. The learning-rate schedule used linear warm-up for the first 10% of optimization steps, followed by cosine decay. Dropout of 0.1 was applied in the ModernBERT attention and MLP blocks and in the classification head.

Training used gradient accumulation to maintain a fixed effective batch size under GPU memory constraints. In the default training script, the per-GPU batch size was 4 windows, and the target effective batch size was 384 windows per optimization step; the number of accumulation steps was computed from the number of GPUs used. This aggregation was used to stabilize optimization for sparse genomic labels and class-imbalanced token classes.

The objective was token-level cross-entropy over *C* = 17 labels, computed only on supervised positions. Let *z*_*i,e*_ be the logit for class *c* at token *i, y*_*i*_ the corresponding label, and *m*_*i*_ ∈ {0, 1} a mask indicating supervised tokens. With class weight 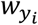, the loss was

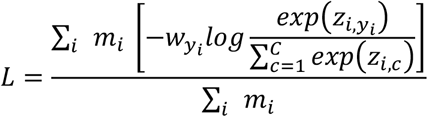

Class weights were computed from the training set using clipped inverse-square-root class frequencies and normalized across classes. Training was monitored using token-level F1 and interval-level precision, recall and F1 for gene-structure features on validation windows. We did not use early stopping.

#### Chromosome-Wide Conditional Random Field (CRF)

During inference, per-base token logits were first aggregated into feature-level logits for five annotation states, namely intergenic, intron, 5′ UTR, CDS and 3′ UTR. For each chromosome and strand, these feature logits were ordered by genomic position and decoded using a fixed-transition Viterbi algorithm. The transition matrices encoded biologically plausible feature order by assigning zero or near-zero probability to disallowed state changes, thereby reducing isolated or structurally inconsistent predictions. Separate transition matrices were used for plant and animal models to reflect domain-specific gene-structure statistics. For animal predictions, an optional CDS-aware Viterbi decoder further incorporated coding-frame and splice-site constraints. The decoded per-base state sequence was collapsed into genomic intervals and exported as GFF3 gene models.

### Post-Prediction Refinement and Validation

After chromosome-scale decoding, predicted GFF3 records were subjected to lightweight structural and protein-level refinement. We first removed very short features and gene models and retained only gene structures passing basic validity checks. For animal predictions, an optional split-gene merge step was applied to adjacent CDS-bearing transcripts on the same chromosome and strand. Candidate pairs were considered only when the intervening gap was within a predefined range and contained compatible splice donor and acceptor motifs. This step was designed to recover gene models that had been fragmented across long intronic or low-confidence intergenic regions.

We then applied ReelProtein as a protein-level plausibility screen. Candidate coding sequences were translated only when they formed complete open reading frames with an ATG start codon, an in-frame TAA, TAG or TGA stop codon, and no internal stop codons. For candidate merges, the procedure allowed small CDS boundary corrections by testing 1 or 2 bp offsets only at near-boundary canonical splice junctions. Protein sequences passing these ORF checks were embedded with a frozen ProtT5-XL-UniRef50 encoder and scored using an ensemble of XGBoost classifiers loaded from Hugging Face. In the default configuration, we used plant-trained ReelProtein classifiers, while the framework allows the scoring model to be replaced for other lineages. Candidate merged models with a mean plausibility score of at least 0.5 were retained. Unmerged gene models were preserved by default unless strict filtering was explicitly enabled. The final refined predictions were written in GFF3 format with updated parent-child relationships for merged models.

### Performance Evaluation Metrics

GeneCAD predictions were exported as GFF3 files and compared with reference GFF3 or GTF annotations using a unified evaluation script. The evaluation comprised five complementary sections, including CDS-based matching, gffcompare-style structural evaluation, splice-site analysis, BUSCO completeness assessment, and site-level error breakdowns. For references lacking explicit exon records, exon structures were reconstructed by merging CDS, 5′ UTR and 3′ UTR features, enabling consistent comparison with GeneCAD predictions.

CDS-based evaluation ignored UTR annotations and assessed performance at the locus, transcript and CDS-exon levels. A predicted transcript was counted as correct only when its complete set of CDS intervals matched a reference transcript on the same chromosome and strand. Structural evaluation was performed using gffcompare-style metrics over full exon structures, including base-level exon overlap, exact exon matches, exact intron matches, intron-chain matches for multi-exon transcripts, and intron-chain-based locus matches.

Site-level accuracy was quantified separately for translation initiation sites, translation termination sites, splice donor sites and splice acceptor sites using strand-aware genomic coordinates. Splice-site composition was further assessed from the genome FASTA by classifying predicted introns as GT-AG, GC-AG or non-canonical. When genome FASTA files and BUSCO were available, predicted spliced transcript sequences were extracted and evaluated with BUSCO in transcriptome mode using the appropriate lineage dataset. Precision, recall and F1 were computed from pooled true-positive, false-positive and false-negative counts for each species. Cross-species summaries were then obtained from the per-species results.

### Evaluation on curated classical and NLR gene sets

To quantify recall at functionally important and structurally complex loci, we evaluated GeneCAD predictions in Zea mays using two curated reference sets that were not used for model training. The first was a maize classical gene set comprising manually curated high-confidence loci from MaizeGDB [45]. The second was a curated set of nucleotide-binding leucine-rich repeat (NLR) genes, which are often difficult to annotate because of high sequence divergence, frequent clustering and repeat-rich genomic contexts.

Reference annotations for each set were extracted from the maize B73 reference GFF3 and compared with GeneCAD predictions using gffcompare. We summarized matches from the gffcompare reference map using a hierarchical stringency framework. High-confidence recovery was defined as reference transcripts associated with either exact intron-chain matches (=) or predictions fully contained within the reference transcript without splice conflict (c). Additional structurally related predictions were recorded separately, including transcripts sharing splice junctions with the reference (j) and retained-intron or related transcript-structure differences (m and n). Transcript completeness was calculated as the fraction of reference transcripts with at least one = or c match, and gene completeness was calculated as the fraction of reference genes with at least one recovered transcript. Because GeneCAD was not retrained or fine-tuned on these curated validation sets, these analyses provide a targeted zero-shot assessment of its ability to recover biologically important maize loci.

## Results

### GeneCAD framework

Current genome annotation approaches fall into two groups. RNA- and protein-supported pipelines, such as BRAKER3, achieve high accuracy but require extensive experimental data. Sequence-only deep-learning systems, including Helixer, Tiberius, and ANNEVO, operate directly on DNA and scale across species; however, these models often require massive training sets and tend to fragment loci or blur feature boundaries due to insufficient structural constraints (Fig. 1a). GeneCAD bridges these paths by implementing a modular architecture that utilizes a lineage appropriate DNA foundation model to encode evolutionary conservation across angiosperms, then fine-tunes a ModernBERT head with a chromosome wide conditional random field to produce coherent, single-transcript gene models (Fig. 1b).

First, we tile assemblies into 8,192 bp windows with a 50% stride to provide localized sequence context for the foundation model’s nucleotide-resolution embeddings. In this study, we use PlantCAD2 to capture plant-specific evolutionary signals. A ModernBERT head processes these embeddings through three parallel feature streams, each optimized for a distinct biological signal: transcriptional boundaries, splice junctions, and coding frames. A gated multilayer perceptron (MLP) subsequently integrates these streams, leveraging the gating mechanism to dynamically weight the importance of boundaries, junctions, and frames before assigning BILOU labels across 5′ UTR, coding sequence, intron, and 3′ UTR regions. This design sharpens boundaries because splice sites and start/stop codons are governed by short, conserved motifs. By isolating splice and frame signals, the model concentrates probability at these conserved motifs, while the BILOU scheme forces state transitions to occur at single-nucleotide resolution. In contrast, UTRs lack a single consensus motif, so evidence accumulates over longer spans and the fused output varies more gradually in these regions.

Second, to enforce biological coherence, we concatenate windowed outputs in genomic order and decode them with a linear-chain conditional random field whose transition matrix encodes legal feature order and splice-phase continuity. Viterbi decoding reduces illegal transitions and merges evidence into chromosome-consistent transcripts.

Third, to increase functional precision, we apply two targeted post-processing steps. To mitigate fragmentation, we reconnect loci split by introns that exceed the 8,192 bp window, provided the resulting sequence supports a single in-frame open reading frame with a consistent splice phase. While this approach effectively recovers the majority of plant gene models, it highlights a technical ceiling for genomes with exceptionally long vertebrate introns. We then remove repeat-driven open reading frames using a protein language-model plausibility score from ReelProtein [9], which increases precision with a small loss of recall. While optimized for plant repeat landscapes here, this filtering logic is adaptable to any protein-coding repertoire.

### Evaluating and curating public genome annotations with masked-motif logistic regression (MMLR)

Genome annotations vary in quality across species and releases, presenting a challenge for supervised models that require high-quality, error-free labels for training. We hypothesized that a sequence-only score could quantify this variation at the transcript level, allowing us to identify and remove poor-quality models while maintaining high diversity in our training data. To address this, we developed the masked-motif logistic regression (MMLR) score, which evaluates each transcript directly from the DNA sequence. MMLR is derived from the underlying foundation model and measures the recoverability of conserved coding and splice motifs from their local context at translation start/stop codons and splice donor/acceptor sites (Fig. 2a). The foundation model is pretrained to predict masked nucleotides from surrounding sequence; therefore, high recovery probabilities suggest that a transcript’s sequence patterns align with features that are frequent and conserved across species. We interpret higher MMLR values as evidence of a biologically credible transcript. Transcription start and termination sites were excluded from this score because these boundaries are often diffuse and lack a single consensus motif in plants (Methods).

We applied MMLR to all transcripts in public annotations for four model plants to validate this curation logic (Fig. 2b). In *Arabidopsis thaliana*, peptide-supported transcripts (n=18,177) achieved a mean MMLR of 0.96, compared to 0.83 for those without peptide support (n=30,279). In *Zea mays*, the means were 0.86 for 27,713 peptide-supported transcripts and 0.69 for 44,826 without peptide evidence. *Solanum lycopersicum* showed 0.92 for 5,390 with support and 0.70 for 28,685 without. *Oryza sativa* showed 0.96 for 3,696 with support and 0.69 for 35,331 without. These contrasts indicate that stronger sequence signals tend to coincide with independent proteomic evidence. The separation is modest, which is expected because peptide catalogs are incomplete; lack of detection often reflects low abundance or tissue-restricted expression rather than poor model quality.

MMLR scoring was subsequently extended to the 65 genomes used for PlantCAD2 pretraining in our primary validation set. Mean transcript-level values ranged from 0.56 in *Ricinus communis* to 0.82 in *Juglans regia*, with a median near 0.73 and an interquartile range of 0.70–0.76, reflecting broad yet interpretable variation across lineages (Supplementary Figure 1; Supplementary Table 1, Fig. 2c).

For GeneCAD fine-tuning, we selected references combining higher mean MMLR, strong assembly contiguity, high BUSCO completeness, and evidence-supported pipelines. Five species met these criteria. Within each, transcripts with MMLR above 0.5 were retained, and the top-scoring isoform was chosen at multi-isoform loci. Retention was consistent across species: *Arabidopsis thaliana* kept 42,323 of 48,456 transcripts (87%), *Glycine max* 66,905 of 80,374 (83%), *Hordeum vulgare* 33,253 of 37,963 (88%), *Oryza sativa* 36,046 of 49,061 (73%), and *Populus trichocarpa* 43,427 of 52,400 (83%). Overall, 60–80% of transcripts were retained, producing a compact and phylogenetically diverse corpus for training and evaluation (Fig. 2c).

### Cross-species benchmarking shows higher accuracy and precision in primary validation clades

To assess performance across diverse evolutionary histories and genomic architectures, we compared GeneCAD to BRAKER3, Helixer, and Tiberius using five held-out angiosperm genomes: *Juglans regia, Coffea arabica, Phaseolus vulgaris, Nicotiana sylvestris* (diploid), and *Nicotiana tabacum* (allotetraploid). These species provide a rigorous cross-species validation set with strong experimental support, enabling evaluation at the nucleotide, exon, and full-transcript levels under exact-match criteria.

GeneCAD consistently outperformed alternative tools across all evaluation metrics (Fig. 3a). Averaged across the test species, the mean F1 at the base level was 0.77 with a range of 0.67 to 0.89, and the exon-level F1 was 0.81 with a range of 0.73 to 0.90. Because UTR annotations in public references remain biologically heterogeneous, we conducted a targeted CDS-only evaluation; under these parameters, GeneCAD achieved a mean CDS-level F1 of 0.47 with a range of 0.37 to 0.64, demonstrating high precision in coding frame resolution. We then applied a highly stringent *gffcompare* style evaluation that requires exact intron chain identity and identical transcript boundaries. GeneCAD maintained robust accuracy in this setting, yielding a mean full-transcript F1 of 0.48 with a range 0.39 to 0.61. To verify the functional viability of these sequence-only predictions, we evaluated the spliced transcript models against lineage-specific BUSCO datasets. GeneCAD recovered an average of 98.5% of complete BUSCOs (range 97.4% to 99.2%), confirming that the predicted structures translate into highly complete gene repertoires. This structural and functional advantage was evident in both diploid and polyploid genomes. For example, *N. tabacum* is particularly challenging to annotate due to duplicated subgenomes and high repeat content. However, the *gffcompare* transcript-level F1 for this allotetraploid still reached 0.42, highlighting the framework’s ability to handle complex genomic architectures (Fig. 3a, Supplementary Table 2).

**Figure 3.**
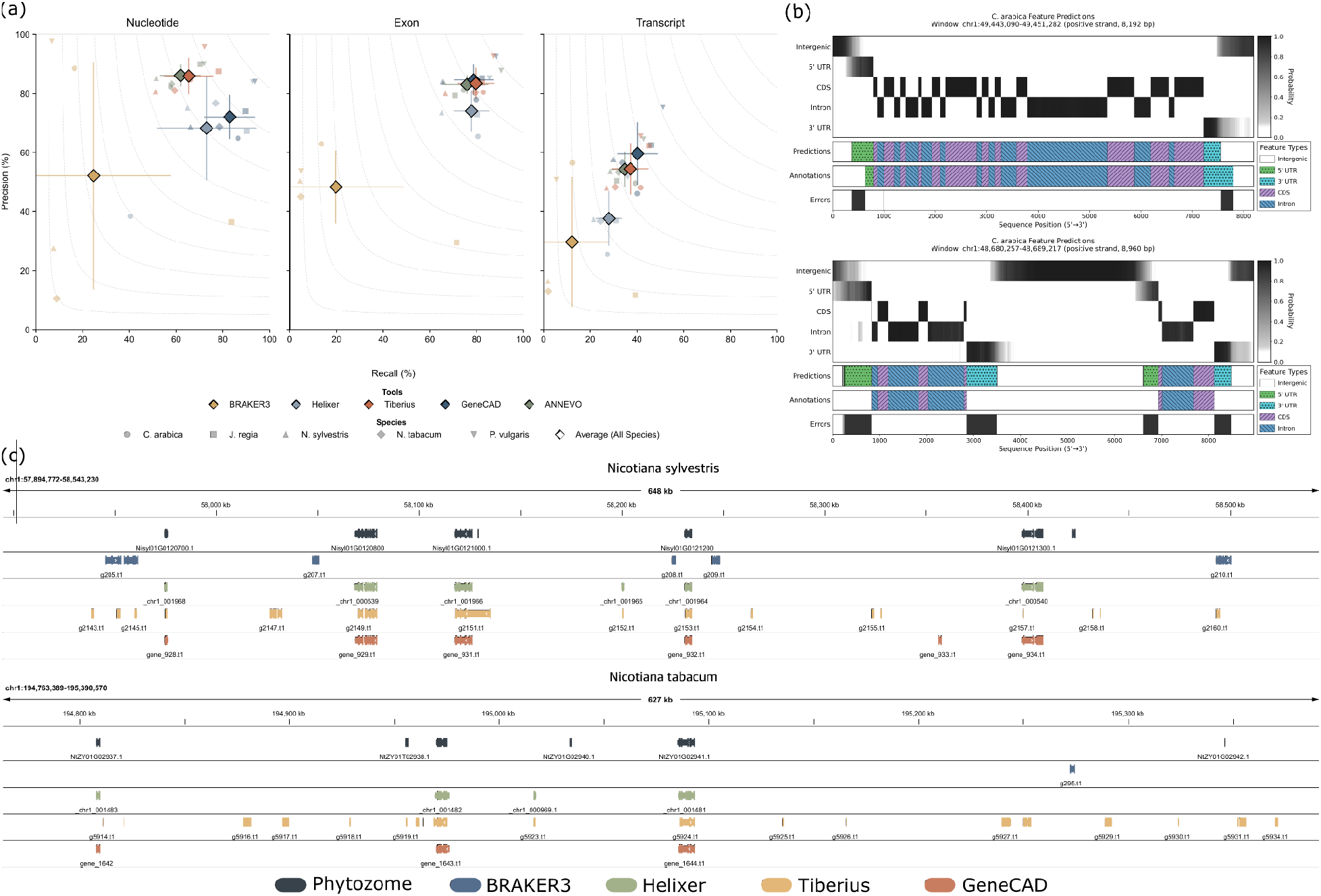
Benchmarking GeneCAD against evidence-supported and sequence-only pipelines in primary validation clades. **(a) Precision–recall across evaluation levels**. Precision–recall plots at nucleotide, exon, and transcript levels compare BRAKER3, Helixer, Tiberius, and GeneCAD on five held-out angiosperms (*Juglans regia, Coffea arabica, Phaseolus vulgaris, Nicotiana sylvestris, Nicotiana tabacum*). Points are colored by species; F1 iso-contours are shown. GeneCAD occupies the best precision– recall region across panels, with the largest gains at the transcript level. **(b) Boundary profiles in an example window**. Per-base class probabilities for GeneCAD are shown above the reference track. Probabilities peak at start/stop codons and splice junctions, while 5′ and 3′ UTRs appear as broader gradients. The error track marks differences relative to the reference. **(c) Representative loci in *Nicotiana***. Browser views in N. sylvestris and N. tabacum compare predictions from Phytozome (reference), BRAKER3, Helixer, Tiberius, and GeneCAD. GeneCAD closely matches curated models and reduces intergenic calls relative to ab initio baselines, including in repeat-rich regions and across subgenomes. The cross-lineage extensibility of these results is further detailed in our animal genome evaluations (Supplementary Note 2).

Boundary profiles showed sharper, higher-confidence transitions at conserved landmarks such as initiation codons and splice junctions across all tools (Fig. 3b, Supplementary Table 2). Predictions across 5′ and 3′ UTRs displayed smoother gradients, which is consistent with the diffuse transcription start and termination boundaries typically observed in plants. In several loci, GeneCAD predicted UTRs not recorded in the reference (Supplementary Figure 2). Although these predictions are counted as false positives under strict benchmarking criteria, they likely represent more complete transcript models than those found in existing annotations.

Visual inspection of representative regions further highlighted differences in specificity (Fig. 3c). Traditional *ab initio* baselines frequently produced multiple gene-like calls in intergenic sequences, often overlapping annotated pseudogenes or transposable-element fragments. By requiring concordant evidence across the transcription, splicing, and translation streams before emitting a gene model, GeneCAD significantly reduced these spurious calls and achieved higher precision at comparable recall. We next tested whether this architectural logic translates to non-plant taxa by adapting the framework to animal lineages. In these vertebrate evaluations, GeneCAD maintained high fidelity at the individual exon level. However, the extreme length of animal introns relative to the model’s context window limited the accuracy of full-transcript assembly (Supplementary Note 2).

### DNA foundation model and curated data drive GeneCAD performance

To understand why GeneCAD outperforms existing tools, we ran ablation tests starting from the full system and sequentially removing each component one at a time. We focused on CDS-level evaluation as the primary metric because UTR annotations in public references remain heterogeneous, and transcription boundaries in our primary validation clade (plants) are diffuse in biology.

Starting from the complete system, we first evaluated the impact of the two targeted post-processing steps. Because inference runs in 8,192 bp windows, some genes with long introns can be split across adjacent segments; reconnecting loci when a single in-frame open reading frame with preserved splice phase is supported restores full-length structures without altering local boundaries. The protein language model screen removes coding-like predictions that lack protein-like regularities. Dropping these refinements reduced precision, with the protein screen yielding a modest gain while recall changed little. Larger declines arose when structure-aware decoding and conservation features were removed. Without the chromosome-scale CRF, feature transitions became incoherent and long loci fragmented. Without lineage-appropriate conservation features, the ModernBERT head failed to learn robust cross-species rules. The modular nature of this requirement is further evidenced in our animal trials, where swapping PlantCAD2 for a mammalian-trained foundation model was sufficient to adapt the architecture to non-plant taxa (Supplementary Note 2). In the plant validation, removing the CRF reduced CDS-level F1 by around 5%, and removing PlantCAD2 reduced it by 30%. Together, these tests show that evolutionary priors from PlantCAD2, structure-aware decoding, and the two refinements provide complementary gains, with PlantCAD2 and the CRF accounting for most of the improvement (Fig. 4a; Supplementary Figure 3; Supplementary Table 2).

**Figure 4.**
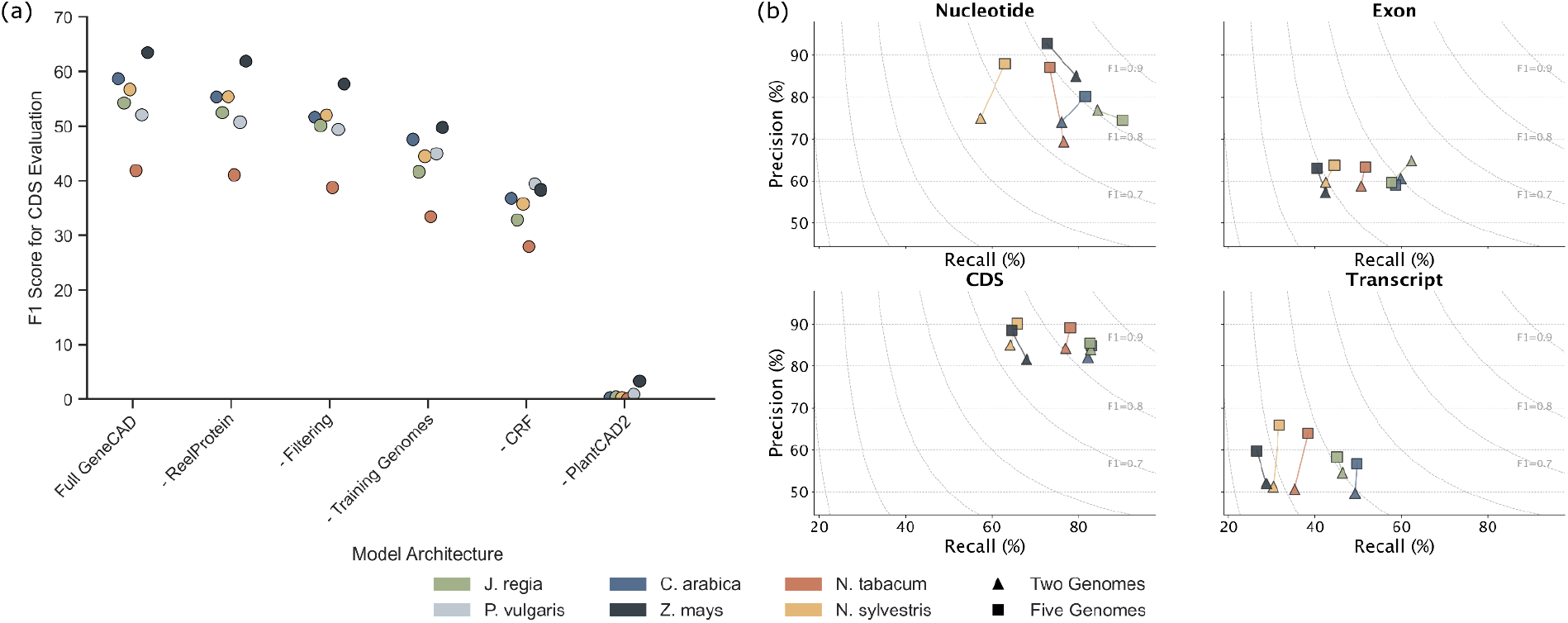
Contributions of modular model components and curated data to GeneCAD performance. **(a) Ablation on CDS accuracy**. CDS F1 scores for each test species are shown as components are removed from the full system (left to right): Full GeneCAD, without the protein plausibility screen (ReelProtein), without post-prediction filtering, trained on two rather than five curated genomes, without the chromosome-wide CRF, without PlantCAD2 Large embeddings, and without PlantCAD2 Small embeddings. Points are colored by species (*Juglans regia, Coffea arabica, Phaseolus vulgaris, Nicotiana tabacum, Nicotiana sylvestris*; *Zea mays* shown for analysis only). Removing lineage-specific foundation model features and the CRF produces the largest drops, indicating that conservation features and structure-aware decoding drive most of the gain. **(b) Precision–recall trade-offs by training diversity**. Precision–recall plots at nucleotide, exon, CDS, and transcript levels compare models trained on two curated genomes (triangles) versus five curated genomes (squares). Curves show F1 iso-contours. Across species, the five-genome model shifts toward higher recall with similar precision, whereas the two-genome model retains most of the accuracy, especially at the CDS level.

We next asked how much curated data the model requires. Starting from the five curated species selected by the MMLR screen, we reduced training diversity from five to two species, retaining *Oryza sativa* and *Arabidopsis thaliana*. CDS-, exon-, and transcript-level F1 values decreased only modestly, and the two-species model retained most of the accuracy of the five-species model. Precision–recall points shifted slightly toward lower recall but followed similar F1 contours, indicating that conservation features transfer well and that a compact, phylogenetically diverse training set is sufficient for high performance (Fig. 4b; Supplementary Table 2). To further evaluate whether GeneCAD generalizes to biologically important and repeat-associated loci, we assessed its predictions in *Zea mays* using two curated gene sets and the B73 transposable-element annotation. Maize was excluded from the GeneCAD and ANNEVO training sets, whereas Helixer and Tiberius included maize during training. This design allowed us to compare maize-excluded generalization with methods that had direct training exposure to maize.

Despite being excluded from the training set, GeneCAD recovered 86.1% (359/417) of classical maize genes with at least one exact or contained transcript match. This represents the highest recovery among all evaluated methods, including those with direct exposure to maize during training (Table 1). It also produced 351 exact classical transcript matches, exceeding Helixer, Tiberius and ANNEVO. For the more structurally complex NLR gene set, Tiberius recovered the largest number of loci under the broader exact-or-contained criterion, whereas GeneCAD produced the highest number of exact NLR transcript matches. Thus, GeneCAD showed strong structural precision for curated loci despite no maize-specific training.

**Table 1.** Curated maize gene recovery and TE-associated coding burden. Gene recovery was defined as the fraction of reference genes with at least one exact or contained transcript match by gffcompare. Exact transcripts denote complete structural agreement with the curated reference transcript. TE-associated coding burden was quantified by intersecting predicted CDS intervals with the B73 transposable-element annotation. CDS bases in TE intervals were calculated after merging overlapping TE intervals to avoid double counting.

| Method | Maize in training | Classical genes recovered | Classical exact transcripts | NLR genes recovered | NLR exact transcripts | TE features overlapped by CDS | CDS bases in TE intervals |
| --- | --- | --- | --- | --- | --- | --- | --- |
| GeneCAD | No | 359 / 417,<br>86.1% | 351 | 65 / 119,<br>54.6% | 52 | 0.44% | 6.08% |
| Helixer | Yes | 276 / 417,<br>66.2% | 274 | 51 / 119,<br>42.9% | 44 | 0.70% | 10.44% |
| Tiberius | Yes | 335 / 417,<br>80.3% | 250 | 83 / 119,<br>69.7% | 35 | 3.67% | 53.91% |
| ANNEVO | No | 281 / 417,<br>67.4% | 218 | 69 / 119,<br>58.0% | 35 | 0.35% | 7.36% |

We next tested whether curated-gene recovery was accompanied by excessive coding predictions in repeat-rich sequence. GeneCAD CDS predictions overlapped 0.44% of annotated TE features, compared with 0.70% for Helixer, 3.67% for Tiberius and 0.35% for ANNEVO. At the base level, 6.08% of GeneCAD CDS bases fell within merged TE intervals, lower than Helixer and Tiberius and comparable to ANNEVO. Together, these results show that GeneCAD generalizes to maize without maize-specific training, combining strong recovery of curated classical genes, high exact-match precision at complex NLR loci and restrained coding prediction in transposable-element regions.

## Discussion

The GeneCAD framework shows that accurate and scalable plant gene annotation is possible without species-matched transcriptomics or proteomics. To our knowledge, GeneCAD is the first framework to successfully combine DNA foundation models with structured decoding to produce coherent, end-to-end transcript models directly from sequence. By coupling pretrained plant DNA, such as PlantCAD2 for angiosperms, with structure-aware decoding and a protein-level plausibility check^9^, this modular system converts evolutionary grammar in sequence into biologically coherent gene models. In our primary validation across held-out angiosperm genomes that include diploids and an allotetraploid, GeneCAD delivers higher accuracy at exon, CDS, and full-transcript levels than current pipelines while using only DNA sequence as its input, improving transcript-level F1 by around 9% relative to Helixer, BRAKER3, and related tools in cross-species benchmarks. While these results establish its efficacy in the challenging genomic landscapes of plants, the underlying architecture is lineage-agnostic. While its application to animal genomes revealed that transcript assembly is currently limited by the 8,192 bp context window, the model successfully maintained structural fidelity at the individual exon level (Supplementary Note 2).

### Toward conservation-driven, sequence-only annotation

The genome sequence contains the signals that specify gene structure, and in principle, gene models can be inferred from sequence alone [29]. Recent models make this practical but highlight distinct trade-offs between evidence-supported and sequence-only approaches. RNA- and protein-supported pipelines, such as the gold-standard BRAKER3 [2] remain highly effective in well-studied species. However, their dependence on species-matched transcriptomics and proteomics limits scalability when experimental data are sparse. To reflect realistic conditions for newly assembled or wild genomes, we used ten RNA-seq libraries per species and no proteomics in our BRAKER3 runs, mirroring the limited-evidence setup described in the tool’s original publication [2]. Under these constraints, BRAKER3 heavily favored precision over recall, particularly in plants. At the nucleotide level, we observed 0.89 precision with 0.16 recall in *Coffea arabica* and 0.98 precision with 0.07 recall in *Phaseolus vulgaris* (Fig 3a). UTRs remained especially difficult to model because UTR training in AUGUSTUS [4] is unstable and not recommended for routine use, which further reduces the number of exact transcript matches.

Sequence-only deep learning offers a complementary path. Helixer [11] and Tiberius [12] operate directly on DNA and avoid assay collection entirely. These tools scale across species by training on massive datasets: Helixer was trained on 186 animal and 51 plant genomes, while Tiberius was trained on 37 mammalian genomes. In our benchmarks, however, these tools often blurred exon and UTR boundaries or fragmented transcripts. This resulted in lower exon- and transcript-level F1 scores relative to GeneCAD, even in cases where base-level performance was similar. Recent work such as SegmentNT [18] extended the foundation-model strategy by fine-tuning the Nucleotide Transformer [19] to segment genic elements at single-nucleotide resolution. The limitation of this approach is that raw probability tracks require extensive heuristic post-processing to generate valid gene models.

GeneCAD builds on this paradigm but bypasses the need for ad hoc heuristics. By decoding lineage-appropriate DNA embeddings through a chromosome-wide conditional random field, GeneCAD yields sharp boundaries and coherent transcripts directly from sequence. When paired with a protein language model filter, this architecture produces fewer intergenic false positives and consistently higher exon, CDS, and full-transcript F1 scores across tested lineages compared to Helixer, Tiberius, and SegmentNT. Unlike Helixer and Tiberius, which train de novo from large curated GFF annotations, GeneCAD fine-tunes a pretrained foundation model on a compact set of only five MMLR-curated plant genomes. Despite this smaller training footprint, the model generalizes effectively to held-out angiosperms that span major clades and include an allotetraploid. At the same time, unlike BRAKER3, GeneCAD retains the practical advantages of sequence-only inference. It requires only DNA sequence, avoids RNA-seq alignment, and still matches or exceeds the accuracy of evidence-supported pipelines in realistic low-coverage regimes while maintaining robustness to genome size, ploidy, and repeat content [10,29]. Finally, while full-transcript assembly in vertebrates remains constrained by inference window limits, the framework successfully identifies individual animal exons. This cross-lineage portability confirms that pairing foundation model embeddings with structure-aware decoding provides a highly flexible framework for genome annotation (Supplementary Note 2).

### Quality-aware learning to improve annotation

Even for well-studied species, reliable gene models coexist with uncertain isoforms, and ambiguity regarding the representative transcript can blur boundaries and inflate intergenic false positives. To address this, we developed a sequence-based confidence metric called the masked-motif logistic regression (MMLR) score. This score allows us to select high-quality transcripts across phylogenetically diverse lineages. In our primary validation set, MMLR uses PlantCAD2 to assess conservation signals in plants (Fig. 2a). On average, the MMLR score correlates well with independent peptide support [10]. However, it also preserves plausible gene models that proteomics frequently under-samples, such as low-abundance [30] or tissue-restricted transcripts [31] and many membrane proteins [32] (Fig. 2b). This curation logic is fundamentally foundation-model agnostic and can be readily adapted to animal clades by leveraging lineage-appropriate DNA representations (Supplementary Note 2).

Curation with MMLR yielded a smaller but cleaner corpus and improved overall structural fidelity. We observed the largest gains at the CDS and exon levels, alongside a reduction in spurious intergenic calls (Fig. 3b). Retention rates remained stable across representative plant assemblies. For example, *Arabidopsis thaliana* retained 87% of transcripts (42,323 of 48,456), *Zea mays* retained 75% (54,383 of 72,539), and *Triticum aestivum* retained 82% (109,184 of 132,624) (Supplementary Table 1).In maize, where pangenome classification is available, the 30,164 MMLR-curated genes cover 85% of the core set (21,046 of 24,716), 72% of the near-core set (1,648 of 2,286), 44% of the dispensable set (2,951 of 6,732), and 42% of private genes (152 of 366) [33]. This indicates strong coverage of conserved loci with the appropriate exclusion of lineage-specific or noisy models. These results demonstrate that conservation-aware representations combined with quality-aware curation provide exceptionally clean supervision for fine-tuning. This approach directly yields sharper boundaries and better splice-phase consistency. While we focus our current claims on curated, high-confidence references, adjusting these thresholds to recover additional, lower-confidence models remains a priority for future work.

### Biological realism and interpretable outputs

Gene annotation is most useful when outputs are interpretable and ready for downstream analysis. Unlike sequence-only models such as SegmentNT [18] that return nucleotide probability tracks requiring substantial post-processing, GeneCAD applies a chromosome-scale CRF to enforce feature order and splice-phase consistency. This design ensures the production of coherent, standard GFF3 gene models without relying on ad hoc heuristics. This structural coherence translates directly to biological utility. In *Zea mays*, GeneCAD recovered a vast majority of the curated classical genes and highly variable NLR disease-resistance loci, maintaining strict structural fidelity across both sets. The robust recovery of NLRs is particularly significant because these genes are frequently embedded in complex, repeat-dense clusters that confound standard sequence predictors. In addition, GeneCAD probability landscapes directly track underlying biological signals. These landscapes peak sharply at initiation codons and splice junctions but vary gradually across 5′ and 3′ UTRs. This pattern is consistent with the diffuse transcription boundaries observed in plant CAGE-seq datasets [34] (Fig. 3b). These probability maps explain several instances scored as false positives under strict matching, suggesting that GeneCAD frequently proposes biologically plausible transcript extensions rather than algorithmic errors.

The integration of a protein language model provides an independent screen for coding plausibility within this modular framework. Using ReelProtein [9], GeneCAD filters predicted coding sequences associated with low plausibility scores, which frequently reflect repeat-derived or nonfunctional open reading frames. Across the benchmark species, this filtering step increased mean CDS precision by 2.8% with less than a one-point loss in recall. For example, CDS precision in *Coffea arabica* increased from 0.67 to 0.74, and in *Nicotiana sylvestris* it rose from 0.82 to 0.86. These refinements yielded transcript-level precision gains of approximately two percentage points overall, producing annotation sets that reduce manual curation effort while preserving sensitivity (Supplementary Figure 3). While these metrics demonstrate the framework’s precision in repeat-rich plant genomes, the underlying protein-plausibility logic remains fundamentally applicable to animal coding repertoires (Supplementary Note 2).

### Limitations and future directions

GeneCAD currently outputs a single representative transcript per locus. While this provides a high-confidence baseline, it inherently misses genuine alternative isoforms and can reduce overall transcript accuracy in regions with highly variable or incomplete UTRs. Additionally, the 8,192 bp inference window limits long-range context (Fig. 3a). As a result, genes containing exceptionally long introns may cross tile boundaries and appear fragmented, although our post-processing logic successfully rejoins most segments when the local splice phase is preserved. Extending this context window through longer-range attention mechanisms and chromosome-scale training will be required to reduce these edge effects. Future iterations of the framework could also score multiple isoforms probabilistically, assigning statistical confidence to alternative splice chains to better capture splicing diversity.

Model supervision can also be made more evidence aware. Rather than relying entirely on fixed labels, the fine-tuning process could incorporate reward functions that favor concordance with orthogonal biological signals, such as splice junction support, full-length isoform assemblies, transcription start and end profiles, ribosome occupancy, and peptide evidence. Preference-based or reinforcement-style optimization would allow the framework to rank competing models without requiring de novo sequencing [35,36]. Training data curation can be broadened as well. Implementing adaptive thresholds for the MMLR score may help retain rare but valid transcripts, and developing lineage-specific protein plausibility models would better accommodate rapidly expanding gene families. These advancements will build naturally on the modular architecture of GeneCAD, moving the system toward complete, isoform-aware annotation at a genomic scale.

The design is not limited to plants. As demonstrated by our successful results in animal clades (Supplementary Note 2), substituting PlantCAD2 with a lineage-appropriate DNA foundation model, while maintaining the same quality-aware curation and structure-aware decoding, enables accurate and scalable structural annotation across distinct evolutionary branches. For established model species, this sequence-only approach significantly reduces the manual curation burden and helps harmonize reference gene sets. For orphan and wild taxa, it provides a reliable, high-fidelity starting point for functional genomics and comparative evolutionary studies. By integrating pretrained DNA representations with strict structural decoding and biologically grounded curation, GeneCAD delivers coherent gene models directly from raw sequence, establishing a robust and generalizable framework for genome annotation across the Tree of Life.

## Acknowledgments

This work was supported by the U.S. Department of Agriculture Agricultural Research Service through a cooperative agreement with Cornell University, the National Science Foundation (grant no. 2240888), and the National Institutes of Health (grant no. R35GM151348 to M.P.). We gratefully acknowledge the SCINet project, the Texas Advanced Computing Center at The University of Texas at Austin (#MCB24097), and the BioHPC Cloud provided by the Bioinformatics Facility at Cornell University (RRID: SCR_021757) for the computational resources that enabled large-scale experimentation. We also thank Lambda (lambda.ai) in partnership with the Open Athena AI Foundation, Inc. for the generous GPU infrastructure grant that supported the validation, containerization, and deployment of this pipeline for the community.

## Data and Code Availability

The source code for GeneCAD is available at https://github.com/plantcad/genecad. Trained model weights are hosted on Hugging Face at https://huggingface.co/collections/plantcad/genecad, and the associated training and development datasets are available at https://huggingface.co/datasets/plantcad/genecad-dev. The complete masked-motif accuracy scores for all analyzed genomes are provided in the Supplementary Tables.

## Author contributions

Z.-Y. L., J.Z. and E.S.B. designed research; Z.-Y. L., A.B., E.C., E.M developed the framework, implemented the MMLR pipeline, and performed model training and benchmarking. Z.-Y. L., A.B., E.C., E.M., M.S., S.-K. H., M.P., J.Z. and E.S.B. performed research and analyses; Z.-Y. L. wrote the manuscript with all other authors’ suggestions and comments.

