## Supplementary Figure 1 for "GeneCAD: Plant Genome Annotation with a DNA Foundation Model"

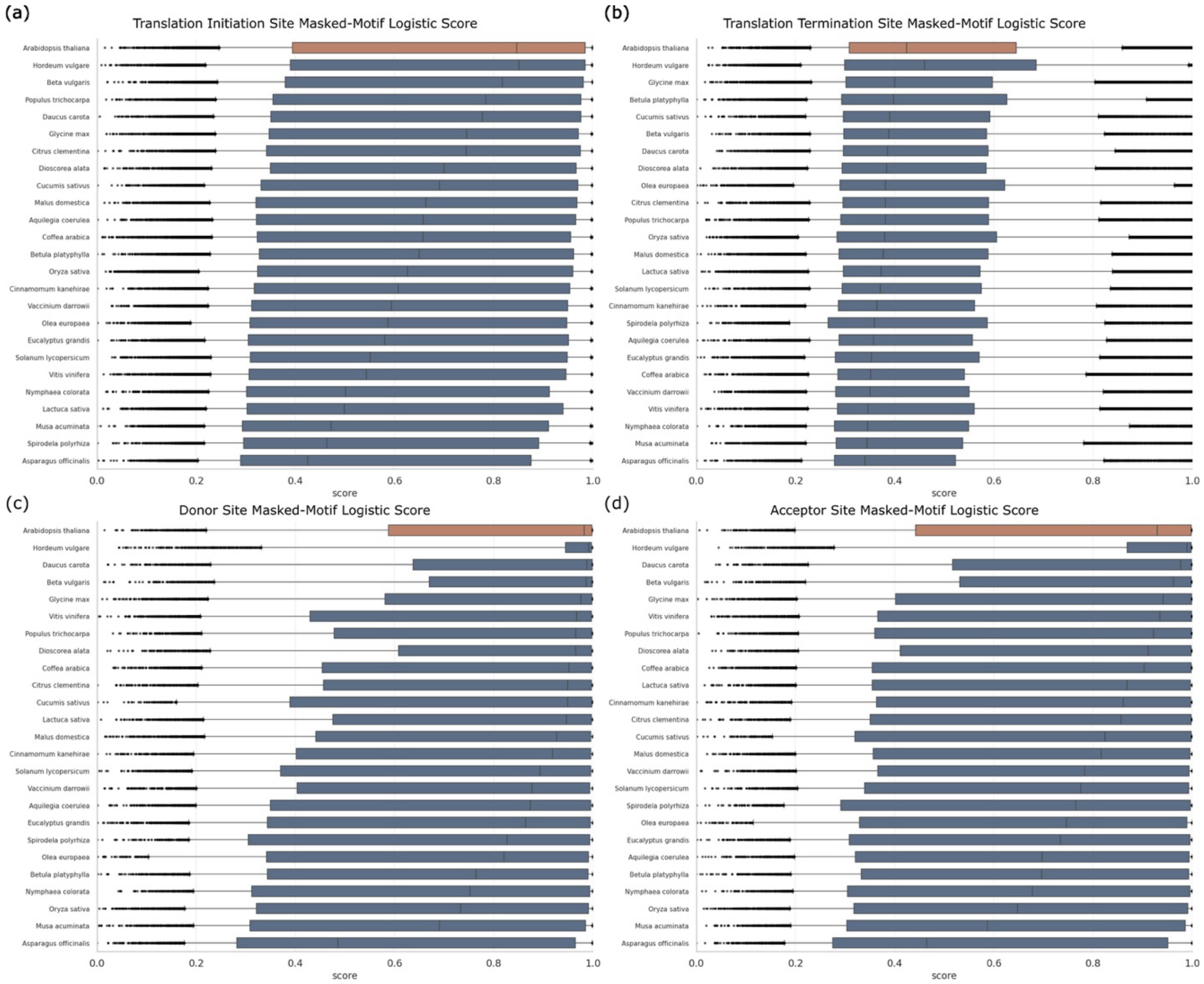


**Supplementary Figure 1 | Species-level masked-motif logistic regression (MMLR) scores for key genomic boundaries**

**a–d**, Distribution of zero-shot PlantCAD2 recovery probabilities for canonical sequence motifs across 24 representative plant genomes. Scores are shown for the **(a)** translation initiation site (ATG), **(b)** translation termination site (TAA/TAG/TGA), **(c)** splice donor site (GT), and **(d)** splice acceptor site (AG). Each bar represents the mean masked-motif logistic regression (MMLR) score per species, with individual transcript values plotted as dots. Species are ordered by mean score. Arabidopsis thaliana and Hordeum vulgare are highlighted to illustrate within-clade variation. Higher scores correspond to motifs that are more consistently recoverable from local context, reflecting stronger conservation at coding and splice boundaries.
