## Supplementary Figure 2 for "GeneCAD: Plant Genome Annotation with a DNA Foundation Model"

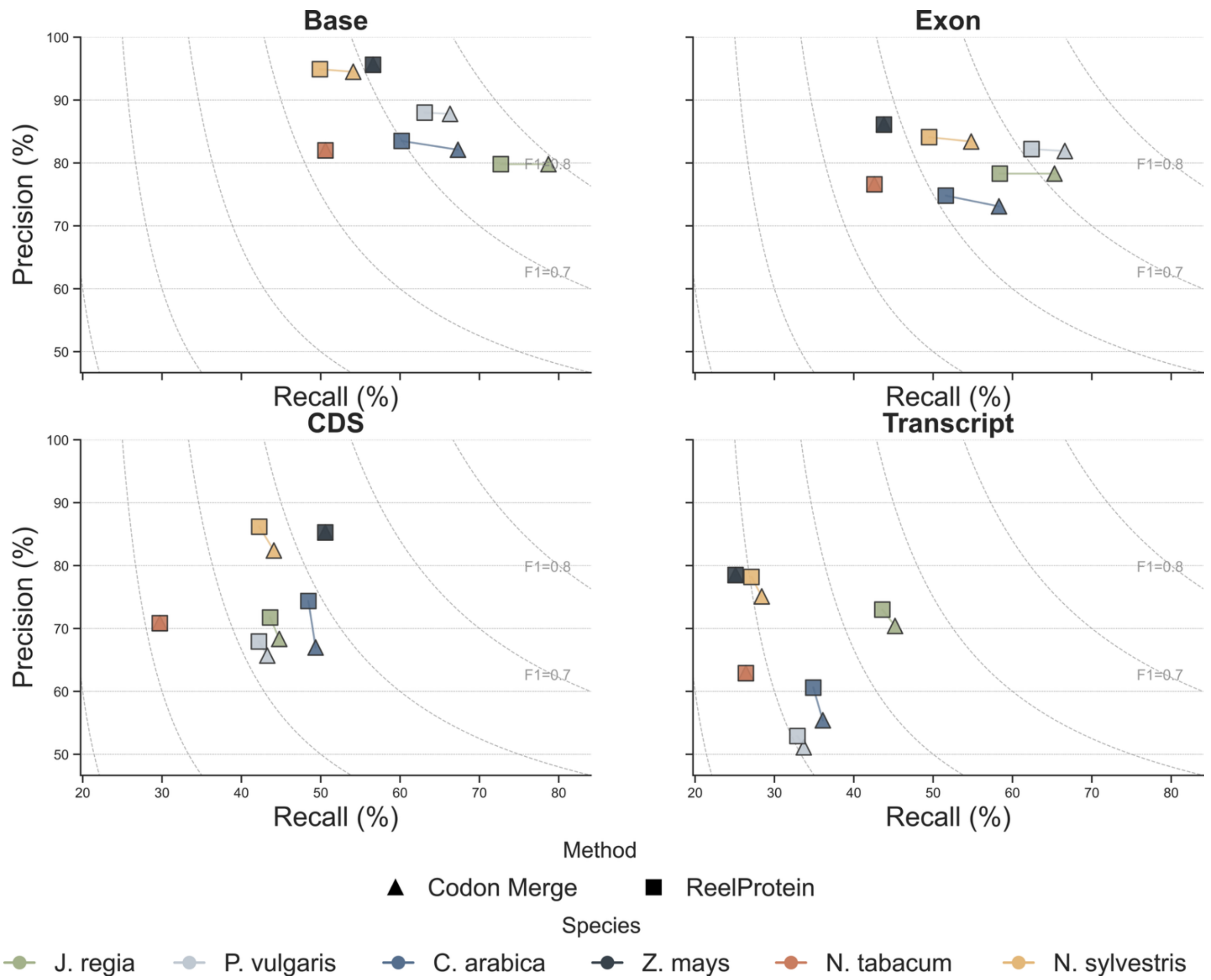


**Supplementary Figure 2 | Effects of codon merging and protein plausibility filtering on precision–recall performance**

Precision–recall plots for GeneCAD predictions across five held-out species (Juglans regia, Coffea arabica, Phaseolus vulgaris, Nicotiana sylvestris, Nicotiana tabacum), with Zea mays shown for reference. Triangles represent models with the codon-merge step only, and squares represent models with the additional ReelProtein plausibility screen. Panels show performance at nucleotide, exon, CDS, and transcript levels. F1 iso-contours are drawn for reference. The protein plausibility filter increases precision with minimal loss of recall, while codon merging primarily improves recall by reconnecting long introns split across inference windows.
