## Supplementary Note 1 for "GeneCAD: Plant Genome Annotation with a DNA Foundation Model"

### Supplementary Note 1 | MMLR function

For each transcript we computed four zero-shot PlantCAD2 probabilities at canonical motifs: translation start (ATG), translation stop (TAA/TAG/TGA), splice donor (GT), and splice acceptor (AG). We then fit an L2-regularized logistic regression in a positive–unlabeled setting on *Zea mays* (positives = classical genes; unlabeled = other transcripts). The model uses the log-odds of the four motif probabilities as predictors:

$$MMLR = -1902 + (1257 \times TIS) + (888 \times TTS) + (571 \times donor) + (847 \times acceptor)$$

We used the resulting MMLR score only to rank species and to retain transcripts (threshold > 0.5; one top-scoring isoform per locus). MMLR was not used as a training label for GeneCAD.
