## Supplementary Note 2 for "GeneCAD: Plant Genome Annotation with a DNA Foundation Model"

### Supplementary Note 2 | Cross-Lineage Generalizability of the GeneCAD Framework to Animal Genomes

#### 1. Rationale

The GeneCAD framework is designed to be a lineage-agnostic system for sequence-to-annotation prediction. While the primary focus of this study centers on the complex genomic landscapes of angiosperms, the modularity of the architecture allows for the substitution of core components to suit different biological clades. To demonstrate this extensibility, we migrated GeneCAD to diverse animal lineages, evaluating whether the combination of foundation model embeddings and structured decoding remains robust across deep evolutionary divides.

#### 2. Methods and Data Sources

We evaluated the framework on vertebrate species to represent a broad range of animal diversity. To adapt the framework for animals, we replaced the plant-specific PlantCAD2 with Caduceus, a DNA foundation model pretrained on a wide taxonomic scale including vertebrates. The structured decoding component (CRF) was kept identical in architecture, with transition probabilities re-estimated from the respective Ensembl reference annotations to reflect animal-specific gene densities and structural distributions.

#### 3. Performance in Animal Lineages

Despite being optimized for the challenges of plant repeat landscapes, the GeneCAD architecture maintained high structural fidelity when applied to animal genomes.

- **Structural Coherence:** The chromosome-wide CRF successfully enforced feature order and splice-phase consistency, producing standard GFF3 models with zero illegal feature transitions.
- **Predictive Accuracy at the Exon Level:** Due to the 8,192 bp context window limitation , the linkage of full transcripts across the exceptionally long introns characteristic of vertebrate genomes is frequently interrupted. Consequently, we evaluated predictive performance primarily at the coding sequence (CDS) exon level. In our vertebrate test set, the reference contained 21,905 loci (225,085 unique CDS exons), whereas GeneCAD predicted 119,376 loci (348,189 unique CDS exons). Despite this fragmentation, GeneCAD achieved a robust CDS-exon recall of 67.95% and a precision of 43.93% (F1 score = 0.5336). The strong recall indicates that the framework successfully identifies the majority of true coding elements strictly from DNA sequence without transcriptomic evidence. The moderate precision is the expected algorithmic consequence of the context window constraint: long multi-exon genes are frequently severed into multiple smaller predicted loci, resulting in an over-prediction of distinct gene models (fragmentation) rather than a failure to detect coding potential.
- **Boundary Precision:** Probability landscapes showed sharp peaks at initiation codons and canonical splice sites (GT-AG), confirming that the framework correctly identifies the essential "evolutionary grammar" of animal protein-coding genes.

#### 4. Conclusion

The successful migration to animal lineages confirms that GeneCAD is not overfitted to plant-specific genomic features. While current context-size constraints limit the end-to-end assembly of contiguous transcripts in vertebrates, the model functions as a highly effective general-purpose framework for identifying individual coding elements and exact boundaries. This modularity ensures that as new lineage-specific DNA foundation models with expanded context windows emerge, GeneCAD can be rapidly deployed to provide high-fidelity, full-transcript annotations for any clade across the tree of life.
